# Harnessing the lymphocyte meta-phenotype to optimize adoptive cell therapy

**DOI:** 10.1101/085910

**Authors:** John Mullinax, Cliona O’Farrelly, Jacob G. Scott, Andreas Buttenschön, Asmaa E. Elkenawi, Fadoua El Moustaid, Alexander G. Fletcher, Clemens Grassberger, Eunjung Kim, Andriy Marusyk, Harry L.O. McClelland, Daria Miroshnychenko, Daniel Nichol

## Abstract

There is an urgent need for reliable effective therapy for patients with metastatic sarcoma. Approaches that manipulate the immune system have shown promise for patients with advanced, widely disseminated malignancies. One of these approaches is adoptive cell therapy (ACT), where tumor-infiltrating lymphocytes (TIL) are isolated from the tumor, expanded *ex vivo*, and then transferred back to the patient. This approach has shown great promise in melanoma, leading to an objective response in approximately half of treated patients [14]. Standard protocols involve characterization of TIL populations with respect to adaptive CD4+ and CD8+ T-lymphocytes, but neglect the possible role of the innate lymphoid repertoire. Due to toxicity and the high cost associated with ACT, the IFN-γ release assay is currently used as a proxy to identify suitable TIL isolates for ACT. Efforts in TIL-ACT for sarcoma, which are pre-clinical and pioneered at Moffitt Cancer Center, have shown that only a minority of the TIL cultures show tumor specific activity in ex vivo IFN-γ assays. Surprisingly, internal melanoma trial data reveal a lack of correlation between IFN-γ assay and clinical outcomes, highlighting the need for a more reliable proxy. We hypothesize the existence of a predictable TIL meta-phenotype that leads to optimal tumor response. Here, we describe preliminary efforts to integrate prospective and existing patient data with mathematical models to optimize the TIL meta-phenotype prior to re-injection.

## 1 Introduction

Soft tissue sarcoma was responsible for 11,930 new cases of cancer and 4,870 deaths in 2015 [1]. The primary modality of therapy for these patients is surgical resection but, unfortunately, 50% of patients will recur and progress with distant metastatic disease. Systemic cytotoxic chemotherapy for these patients offers median survival of 14.3 months [11] and until recently targeted therapy improved median overall survival by only 1.8 months when compared to placebo for patients that have progressed on first line systemic chemotherapy [26]. The simple fact that this therapy was tested in randomized fashion against placebo demonstrates the incredible lack of options for patients with metastatic soft tissue sarcoma. In a recent study Tap and colleagues reported and impressive overall survival gain for the combination of doxorubicin with olaratumab, a monoclonal antibody targeting the platelet-derived growth factor receptor [ref]. This promising result is an improvement of the generally poor outcomes in soft tissue sarcoma, though the low increase in progression free survival (2.5 months) and low objective response rate and duration illustrate the difficulties in treating STS and the need for alternative approaches.

Recent advances in immunotherapies, a collective term for drugs that stimulate an immune response and also for cell-based therapies, have led to improvements in outcomes in a number of cancers. Adoptive cell transfer therapy (ACT) with tumor-infiltrating lymphocytes (TIL) involves obtaining a sample of a patient's tumor, followed by the massive expansion *ex vivo* of T cells from the tumor, and the reinfusion of these T cells back into the patient. This approach has resulted in complete and durable responses in metastatic melanoma [17, 18, 19]. The potential to personalize treatment using active immunotherapy with transferred lymphocytes has been identified in multiple tumor types [5, 13, 20, 25]. A high density of immune cell infiltrates such as TIL in human melanoma and other solid tumors has prognostic value [8]. In sarcoma, the immune infiltrate has been correlated with prognosis [20, 21, 25], and other groups have described the importance of TIL in resected specimens [5, 13, 22, 24].

The infiltration of CD3+ cells has been shown to be a positive prognostic indicator for patients with sarcoma 20, and there are reports of adoptive cell therapy for patients with sarcoma in the literature [4, 5]. One trial reported using T cells genetically modified to recognize the cancer-testis antigen NY-ESO in patients with synovial cell sarcoma 16. A more recent report described the administration of chimeric antigen receptor T cells CAR-T) designed to recognize the Her2 receptor in patients with osteosarcoma [2]. Both cases focused on a clonal population of cells expressing a single antigen. This model is less likely to be effective than TIL therapy in patients, as there is significant antigen heterogeneity within tumors, and selection of resistant subpopulations not expressing the target of the chimeric antigen receptors will likely impair the durability of any response.

### 1.1 Preliminary data

Our lab has successfully generated TIL from resected sarcoma specimens [10]. A total of 9 patients were enrolled on a pilot study and TIL were grown from 40 percent (2/5) of liposarcoma specimens, with no TIL grown successfully from other histologies. The phenotype of these sarcoma derived-TIL indicates a significant population of CD8+ T cells, in some cases with proportions similar to those observed in melanoma. There are two significant differences in the sarcoma TIL and melanoma TIL, however. First, there are other significant populations of lymphocytes within the resected tumors, in particular a population of CD3+ lymphocytes that also express CD56 and are believed to represent Natural Killer T-lymphocytes (NKT) cells as seen in Figure 1. Second, the reactivity of the TIL, as measured by IFN-γ release assay, is less than that of melanoma TIL. Despite some limitations, TIL derived from two patients have been expanded using the rapid expansion protocol (REP) previously used for the melanoma clinical trials, with a similar degree of expansion, indicating that the basic methods derived from that experience are applicable to sarcoma.

**Figure 1:**
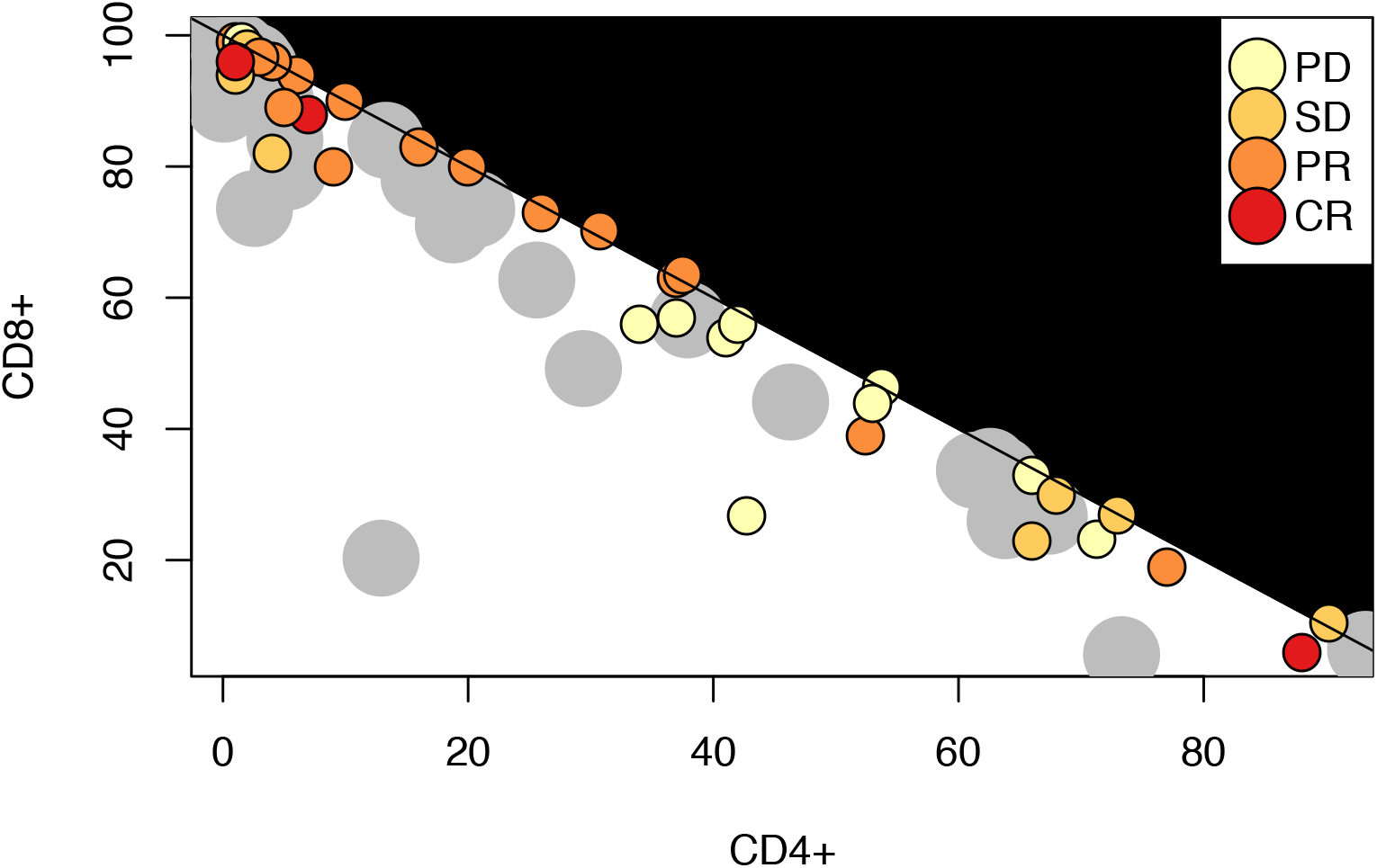
Preliminary data from Mullinax lab suggests that the immuno-phenotype of sarcoma TIL populations (Grey dots) is significantly more heterogeneous than those in melanoma (colored dots). Dots now on the diagonal line represent cells that are neither CD4 or CD8 positive. Colored dots represent clinical response in a trial of melanoma ACT. As data are preliminary for sarcoma, there is no clinical response information.

### 1.2 Rationale for this study

Due to toxicity and the high cost associated with ACT, the IFN-γ release assay is currently used as a proxy to identify suitable TIL isolates. Efforts in TIL-ACT for sarcoma, which are pre-clinical and pioneered at Moffitt Cancer Center, have stalled due to widespread absence of activity in *ex vivo* IFN-γ assays. Surprisingly, internal melanoma trial data reveal a lack of correlation between IFN-γ assay and clinical outcomes, highlighting the need for a more reliable proxy. We hypothesize that there is an optimal balance of lymphocyte populations that will yield clinically significant reactivity to tumor in an *in vitro* assay. The primary purpose of this project is to identify this optimal meta-phenotype of the TIL subpopulations in order to develop a meaningful treatment for patients with advanced sarcoma.

This document represents an initial report of work done at the 5th annual Integrated Mathematical Oncology workshop at the Moffitt Cancer Center. At this workshop, teams of scientists from a variety of backgrounds come together to build mathematical models to answer outstanding questions in oncology. This report represents a suite of mathematical models developed during this workshop (held November 2015). The central tenet around which these models have been built is that for optimal response to a heterogeneous cancer, there must be a concomitant heterogeneous lympoctye population in the ACT. The remainder of this paper is structured as follows. First, we present a simplified Boolean network approach to allow for complexity reduction in the immune response. We then utilize this reduced model of the immune system to create a system of ordinary differential equations to understand the immune cell population balance in general, and the role of checkpoint inhibitors in particular. We end with a simplified model of tumor-immune co-evolution in which we utilize a system of coupled partial differential equations (PDEs) representing the genetic heterogeneity of the tumor and the T-cell population on an abstracted ‘genotype’ space.

## 2 Starting from minimal complexity: a Boolean network approach

The dynamics of human immune response are governed by a complex network of activating and inhibiting interactions both between distinct immune cell types, comprising both adaptive and inert immune respone, and between these immune cells and their microenvironment. Thus, to build a tractable model of immune dynamics it is imperative that we avoid the explosive blow up of the parameter space associated with an attempt to model all of the known components. A such, our modeling begins with a reduction of the complex immune-interaction network to a simplified, qualitatively similar analog. We highlighted five key classes of immune cell type whose interactions are central in driving immune response: NK cells; NKT cells; CD4+ cells; CD8+ cells; and double negative (with respect to CD4, CD8) cells. Each of these classes encompasses a collection of distinct immune cell types that were identified to share qualitatively similar behaviors. Each class was assigned a name corresponding to the most representative cell-type within it.

Intuitively, the state of the immune system can be associated with the abundances of each of the constituent cell-types, with these abundances changing in response to external or environmental stimulus (e.g. infection, emergence of tumor cells). However, the precise dynamics of the immune response, and accurate estimates for the abundances of immune cells in an inactive or active immune state, are difficult to determine. Following the precedent of previous modeling in development [3], cell-cycle regulation in yeast [6] and mammalian cells [7] we forgo modeling the precise immune cell dynamics and focus instead on the state of each immune cell population as active, corresponding to a high abundance, or inactive, corresponding to a low abundance. This simplification reduces the dynamics to that of a Boolean network model governed solely by the nature of interactions between cell-types, rather than the precise physical parameters governing those interactions.

A Boolean network, first introduced by Kauffman as a mathematical tool to study the dynamics of gene regulatory networks [12], assigns one of two states (the Boolean 0 or 1) to each cell-type indicating whether or not it is upregulated (present in high abundance). Interactions amongst these cell-types are characterized as either *activating* or *inhibiting*, indicating that the upregulation of one cell type promotes (respectively inhibits) the upregulation of another. These interactions are encoded as a matrix M ∈ {−1, 0, 1}^*K × K*^ where *K* denotes the number of interacting objects (in our case *K* = 5 immune cell classes), a 1 indicates upregulation, a −1 inhibition and a 0 no interaction. This discrete, parameter-free, encoding of the interactions allows us to exhaustively explore the space of potential interactions amongst cell-types, identifying those that provides reasonable descriptions of empirical observations, and deriving further simplified models that capture the dynamics of larger systems.

A Boolean network determines how the system of immune classes evolves over discrete time steps. Over a single time step, the state (active or inactive) of each immune class is governed by the update rule

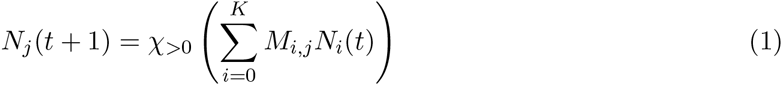

where *N*_*j*_(*t*) ∈ {0, 1} indicates the state of the *j*^th^ cell-type at time step *t* ∈ ℕ. The initial conditions, *N*(0), for these dynamics are chosen dependent on the underlying process that is being modeled.

To identify potentially suitable interaction models for the immune system, we considered three sets of initial conditions and simulated the dynamics of the network. From these dynamics we classified a candidate immune interaction model as suitable or unsuitable. The three suitability tests are as follows. In the first, all cell types in the initial conditions are set as inactive as we propose that, in the absence of external stimulation, each immune cell-type should remain inactive. All networks for which the entirely inactive state is a fixed point (under the update rule of Equation 1) passed this test. In the second, an initial condition was taken as each cell-type activated. We proposed that an active immune system, in the absence of external activating stimuli, should return to an inactive state. In the final suitability test, the initial conditions were taken to correspond to an entirely inactive immune system. We extended the network model to allow constitutive activation of an immune cell-type, simulating an external stimulus warranting an immune response (See Figure 2(C)). We classified a network as suitable under this test is the initially inactive immune system becomes fully active in the presence of the external stimulus.

**Figure 2:**
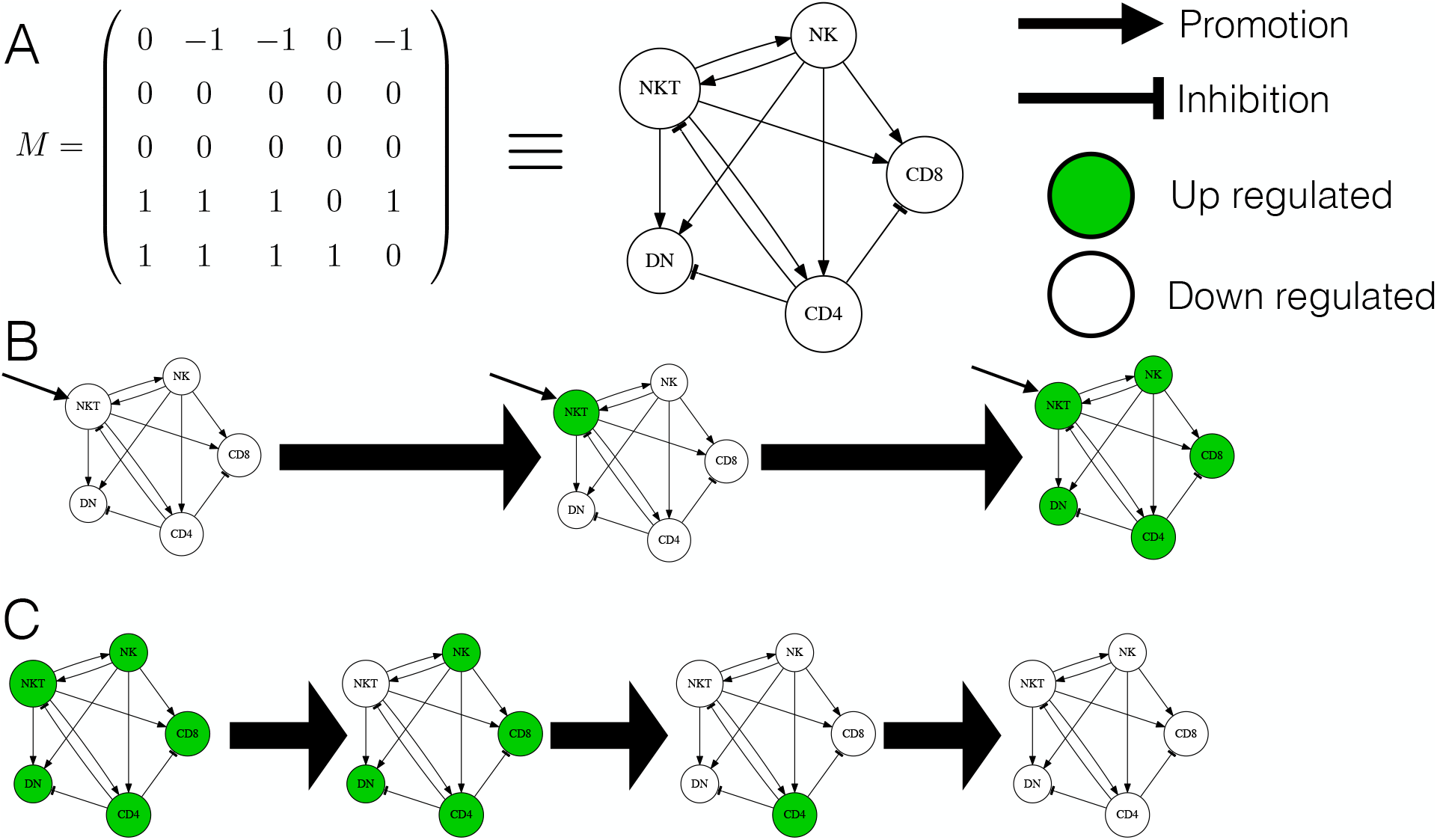
The Boolean interaction network. A. The interaction network and associated matrix M. B. External stimulation “switches on” the network, causing up-regulation of all cell types whilst the external signal persists. C. In the absence of external stimulation, all cells return to a homeostatic down regulated state.

Amongst the potentially suitable networks found through this method, Figure 2 shows the suitable interaction network between the five cells that is most consistent with presently accepted knowledge of immune dynamics. Note that, by construction, this network exhibits a homeostatic property that is vital in immune modelling: a stable ‘off’ state; the ability to ‘turn on’ in the presence of external stimulation (Figure 2(b)); and the ability to ‘turn off’ when this external stimulus is removed.

Despite the reduction in the space of parameters associated with the construction of the Boolean immune interaction model, the number of components is still too high to allow a tractable model for the more precise dynamics of the immune-cell abundances to be constructed. For the five-type immune interaction model derived above, as many as 60 physical parameters may need to be derived.

To further simplify the immune interaction network, and in preparation to build a more precise ODE model, we identified dependencies within the five-type interaction network that permit the grouping of immune cell-types according to their functional role instead of their phenotypic behaviors. Specifically, we noted that the double negative (DN) cells have no outgoing regulatory connections which can be removed (formally combined with the NK/NKT class which determine the regulatory status of DN). Further, the NK and NKT cell types exhibit reciprocal activation and can thus be combined without effecting the dynamics of other immune cell-types. This network reduction is shown in Figure 3. The resulting network exhibits the same homeostatic properties of our biologically derived network (Figure 2), but with a substantially reduced parameter space.

**Figure 3:**
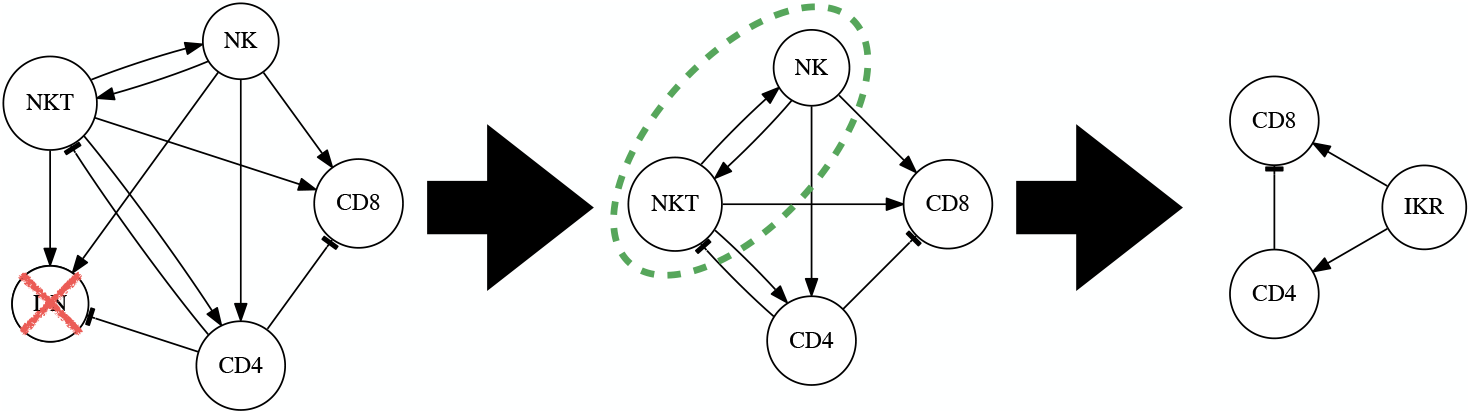
The reduced immune interaction network. Double negative (DN), natural killer (NK) and natural killer T cells (NKT) are grouped by their regulatory function as “innate killer” (IKR) cells.

The process of derivation and analysis of an immune interaction model through Boolean network analysis systematically determines a suitable model. Critically, this process has allows us to drastically reduce the number of parameters that must be estimated to produce a more precise model of immune response; as we will in the following section. Further, by exhaustively exploring the space of potential interaction networks and identifying those that possess critical properties of immune response (e.g. homeostasis and activation), we have derived a tool for rapid ‘sanity checks’ of hypotheses for immune cell interactions. Such a modelling approach widely applicable could prove valuable to other models of immune dynamics, or to other modelling of complex dynamics systems in general.

## 3 Quantifying the dynamics of the immune balance

The Boolean network model is an excellent representation of interactions among the immune cells. Although this model does not incorporate a significant biological complexity, it is able to inform us which interactions are most stable to perturbations. We used this information to identify the main immune cell categories. Figure 4A shows the interactions between tumor cells and three immune cells compartments from the Boolean network model. Each cell population can grow at some rate. Both CD8+ (E) and innate killer (N) cells inhibit tumor growth. Innate killer cells stimulate both CD8+ and CD4+ (R) cells. Finally, CD4+ cells inhibit activity of CD8+ cells. The interactions are described by a system of ordinary differential equations (ODEs) given as

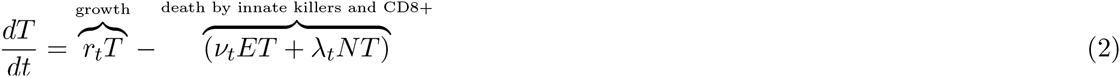

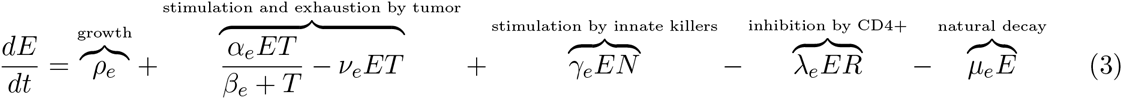

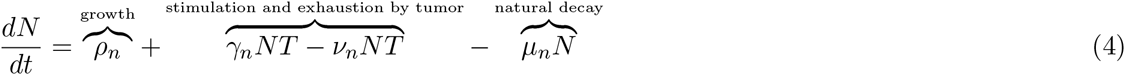

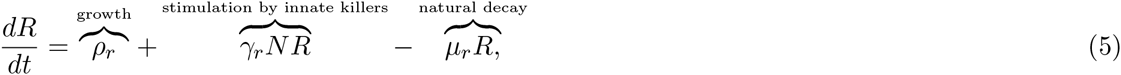

where *T* represents tumor cells, *E* represents CD8^+^ cells, *N* represents innate killer cells, and *R* represents CD4^+^ cells.

**Figure 4:**
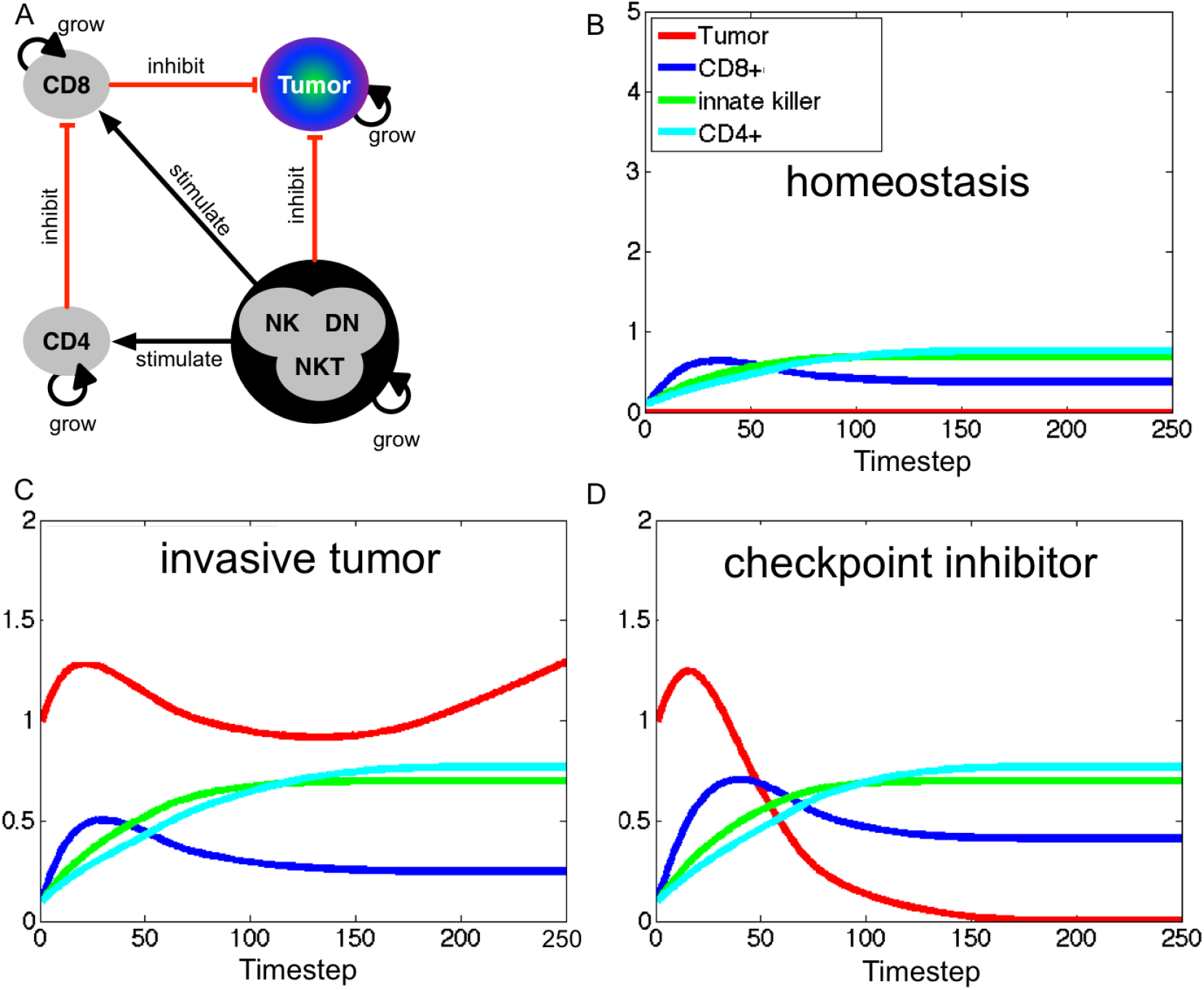
A. Schematic of a compartmental model composed of four compartments, tumor, CD8+, CD4 and Innate Killer (NK,DN,NKT cells). B. A homeostatic state of CD8+ (blue), CD4+ (cyan), and innate killer cells (green) C. A representative case of invasive tumor where immune cells failed to control tumor growth (red) D. A representative case of effect of checkpoint inhibitor.

The parameter values were set to be *r*_*t*_ = 0.34, *v*_*t*_ = 0.5, *λ*_*t*_ = 0.3, *ρ*_*e*_ = 0.3, *α*_*e*_ = 0.31, *β*_*e*_ = 20, *v*_***e***_ = 0.19, γ_**e**_ = 0.1, λ_e_ = 2.0, *μ*_*e*_ = 0.01, *ρ*_*n*_ = 0.14, *γ*_*n*_ = 0.02, *v*_*n*_ = 0.5, *μ*_*n*_ = 0.1, *ρ*_*r*_ = 0.1, *γ*_*r*_ = 0.1, and *μ*_*r*_ = 0.2. Initial conditions are set to be *T*(0) = 1, *E*(0) = 0.1, *N*(0) = 0.1, and *R*(0) = 0.1. The model recapitulated immune homeostasis by setting *r*_*t*_ = 0 and *T*(0) = 0 (Figure 4B). Three immune cell compartments find their stable states just after timestep 50. An unsucessful immune surveillance of tumors was also simulated (Figure 4C). As the tumor compartment grows (red), more CD+8 cells (blue) and innate cells (green) were recruited and expanded. The recruited immune cells were able to reduce tumor growth initially. As CD8+ cells grow, CD4+ cells were expanded (cyan), resulting in the reduction of CD8+ cells and hence increase of tumor volume (rebound of red line at timestep 100).

We also utilize this model to predict effects of checkpoint inhibitors on tumor growth. Checkpoint inhibitor therapy is based on removing inhibitory pathways blocking T cells responses [9]. This therapy can be performed using antibodies that help decreasing CD4+ T cells down-regulation of CD8+ T cells. The therapy has been used on patients with melanoma and has in fact lead to regression of tumor growth [15]. In order to illustrate the effect of checkpoint therapy, we decreased the parameter λ_*e*_ by 50%, and show that it was enough to eradicate the tumor (Figure 4D).

The ODE model provides qualitative information on how tumor and heterogeneous immune population interact with each other, and can be used to explore the effects of different therapies aimed at enhancing the immune response, for example checkpoint inhibitor. While considering a heterogeneous immune population, we assume the tumor to be homogeneous in this example. To address the effect of intra-tumor heterogeneity on immunotherapy outcome, we develop another mathematical model in the following section.

## 4 Understanding T cell and tumor co-evolution and heterogeneity

In the previous two models, our focus was on the heterogeneity in the TIL population, while tumor cells were modeled as a homogeneous population. However, many tumors are themselves highly heterogeneous, as shown in many recent multi-sample sequencing studies [23].To address this additional complexity, we develop a mathematical model to investigate the effect of intra-tumor heterogeneity on ACT treatment outcome. Here, we define ‘treatment outcome’ by the interval of disease-free survival, i.e. the duration that the total cancer cell population remains below a given threshold value following treatment. For this reason we choose cytotoxic (CD8) T cells, which can eliminate cancer cells, as the representatives of the immune system in our model.

We denote the cancer cell population by *C*(*x, t*) and the cytotoxic T cell population by *T*(*x, t*), where *t* ∈ ℝ^+^ is time and *x* ∈ [0,1] denotes an abstract cell genotype. The abstract genotype is discussed in detail later. The model equations are given by

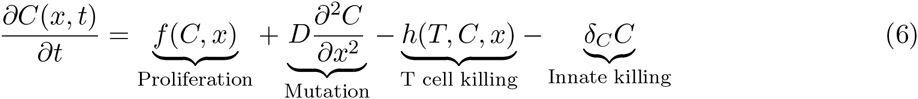

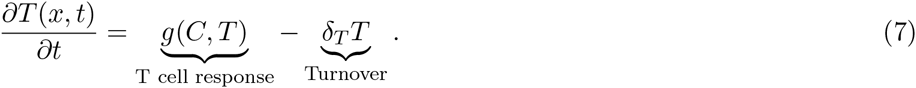

We make several assumptions about the underlying biology to arrive at this model. T-cells detect their targets by constantly sampling the antigens presented on all cell surfaces via the major histocompatibility complex (MHC). Cytotoxic activity only occurs if a match is found. We capture this process by requiring that the genotype *x* of the cancer cell and that of the T-cell match exactly. Further, we assume that the cancer cell genotype undergoes small successive mutations, either due to epigenetic or genetic changes. These mutations are modeled using a diffusion process in the abstract ‘genotype space’. Cancer cells grow using a logistic growth law *f* (*C, x*). Cancer cells are eliminated in two different ways. The innate immune system provides a small constant elimination rate of cancer cells given by *δ*_*C*_. Cytotoxic T cell killing is described by function *h*(*T, C, x*). T-cells are constantly produced and selected for in the thymus, we thus assume that all T cell phenotypes are available at low numbers. Once the T cells recognize their targets they undergo rapid expansion. Both of these effects are included in *g*(*C, T, x*).

The final ingredient of this model is a fitness landscape in *x*. This landscape represents the balance between cancer cell growth and its ability to evade the immune response. We assume that the growth rate *f* (*C, x*) is a decreasing function in *x*. While the immune cell elimination rate *h*(*T, C, x*) is an increasing function in *x*. Thus genotypes closer to *x* = 0 grow fast, but are less resistant to the immune response. On the other hand genotypes closer to *x* = 1 are more resistant and grow at a lower rate.

With this model we study two scenarios; the development of cancer in the presence of an immune response; and various immunotherapeutic treatment interventions. All treatment interventions resemble ACT. That is, at treatment time we add a given distribution of T-cells to the present T-cell population of the model. As mentioned, treatment outcome is measured using the disease free survival time.

**Figure 5:**
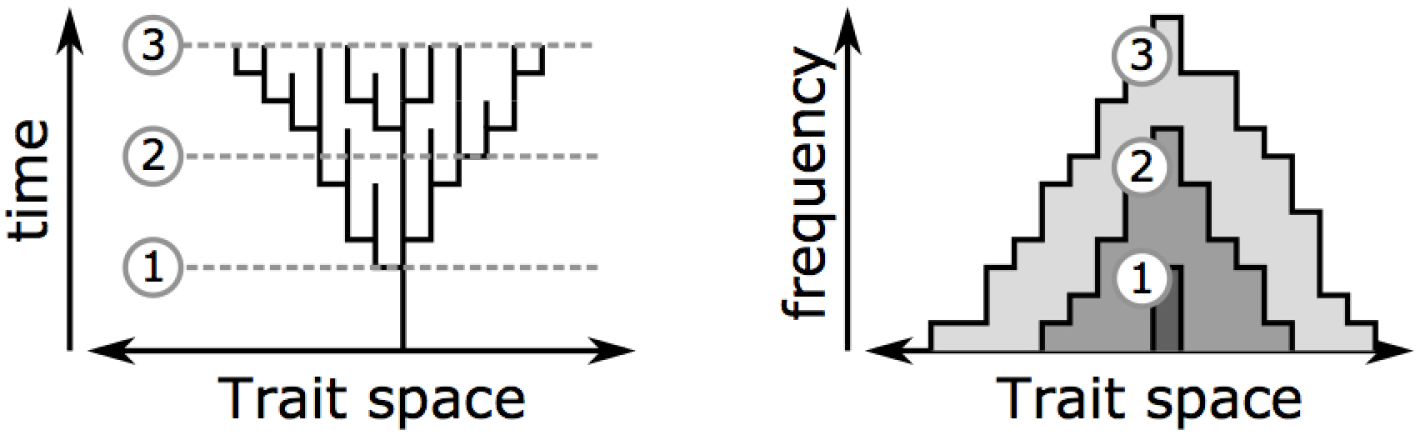
Schematic of the PDE model. We represent the branching evolution of the T-cell and tumor populations as diffusion on a ‘trait-space’.

### Results

In the presence of a fitness landscape, we observed the development of a heterogeneous cancer cell population, characterized by several peaks in phenotype space. Two treatment scenarios are considered. Treatment with all possible T-cell phenotypes and only T-cell phenotypes sampled from the present TIL. The latter matches the adoptive T-cell protocol best. In particular, one can consider cases in which the sampling is not perfect i.e. some sub-clones are not sampled and are missing from the T-cell population. In this case, treatment outcome is poor compared to the constant treatment case, see Figure 6 and Figure 7. The cancer cell populations corresponding to the missing T-cell populations quickly repopulate the tumor site. In conclusion, the model suggests that the treatment of highly heterogeneous cancers is challenging, because the emergence of a resistant sub-clone or a fast outgrowing sub-clone has to be avoided.

**Figure 6:**
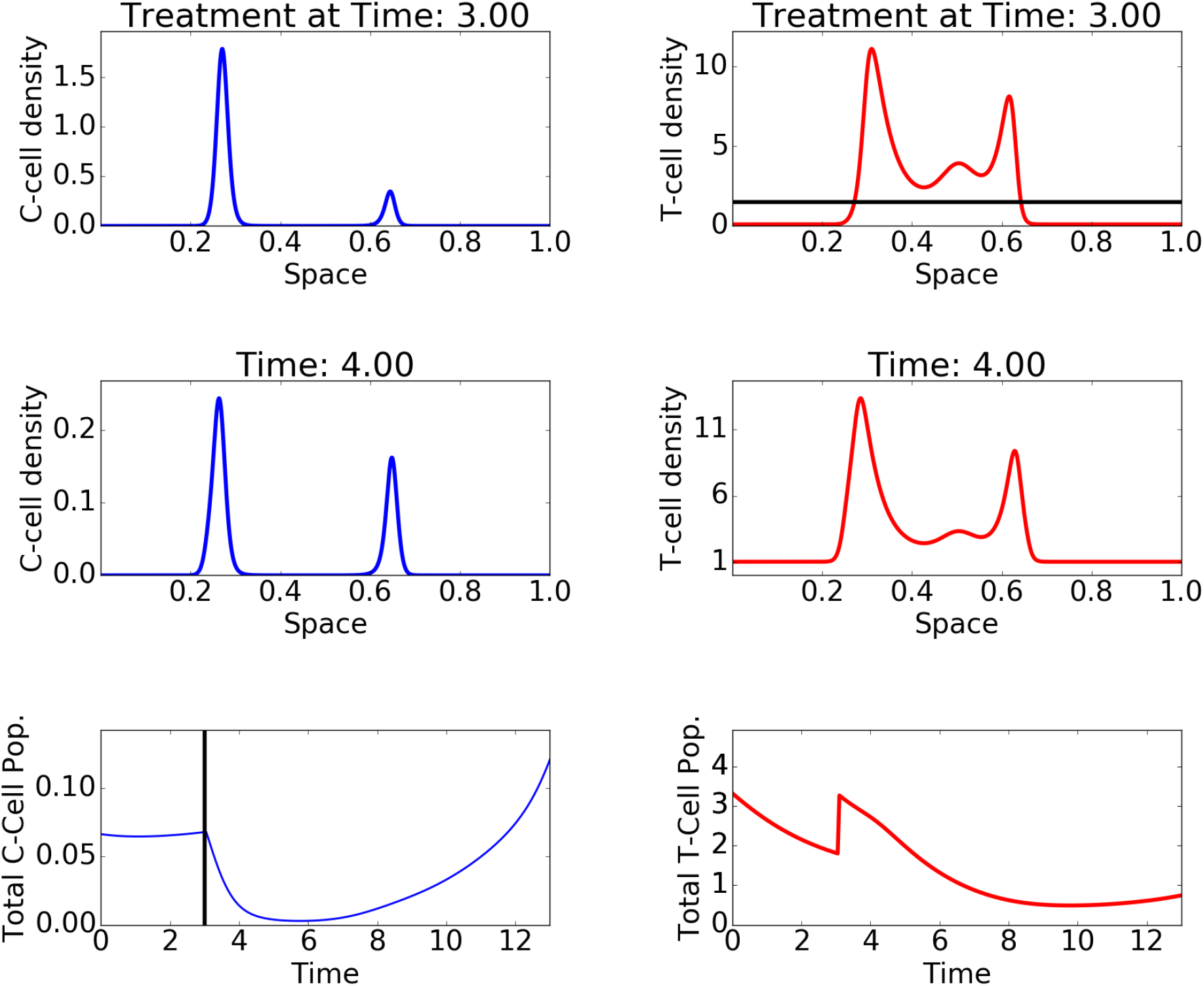
Simulation snapshots of the PDE model. The plots on the left show the evolution of the cancer cell population, while the plots on the right show the T-cell population. The last row of plots shows the evolution of the total cancer cell and T-cell load at the tumour site. The injected T-cell population is depicted in black. Here a constant treatment is given.

**Figure 7:**
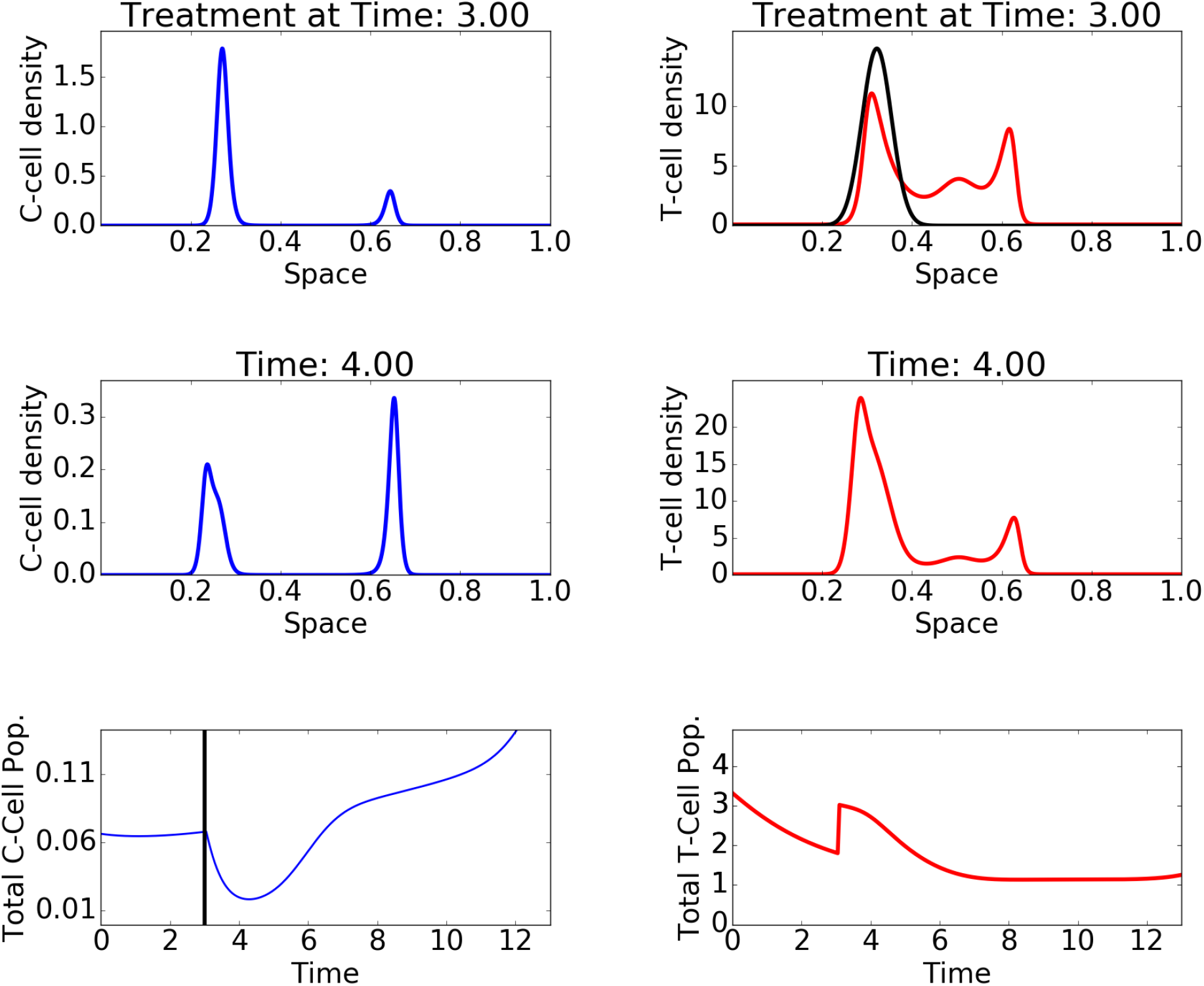
Simulation snapshots of the PDE model. The plots on the left show the evolution of the cancer cell population, while the plots on the right show the T-cell population. The last row of plots shows the evolution of the total cancer cell and T-cell load at the tumour site. The injected T-cell population is depicted in black. Here a sampled (targeted) treatment is given.

## 5 Discussion

During the course of this workshop we generated the hypothesis that, for the treatment of sarcoma with ACT, a heterogeneous population of T-cells is beneficial. This stands apart from the data in melanoma which suggest that only CD8 positive T-cells are meaningful participants in the antitumor immune response. Secondary to this hypothesis, we generated three complementary and increasingly complex mathematical models.

Our Boolean network model provides a methodology to reduce the complexity of the multi-compartment immune system to allow for tractability in what would otherwise a poorly constrained, undetermined problem. We demonstrate that the simplified system can describe the fundamental aspects of the immune response, namely a stable ground state, a response to external stimuli and a return to homeostasis. The reduction in dimensionality is the basis for the next two models and reduces the number of necessary parameters considerably.

Our ODE system suggests several different ways to ‘tip the balance’ of the immune response in sarcoma immunotherapy. The strength of this model is the simplicity inherited from the Boolean network model combined with the interaction of the lymphocyte subsets and the time evolution of the immune response. Given proper patient data to parametrize the model it could yield important insights into optimal timing and dosing strategies for differing immunotherapeutic approaches, such as checkpoint inhibitors.

The PDE model described in the last chapter focuses on the heterogeneity of the tumor cell population, which was not taken into account in the PDE model. The model includes the phenotypic heterogeneity in tumor cells as well as T-cells and demonstrates how depending the response is on a “broad” (in a phenotypic) sense immune response, and how tumors can escape even an effective immune response by evolving in trait space. This can yield important conclusions especially for adoptive cell transfer approaches, in which we can control the relative concentration of cells to be re-injected and tailor it to the lymphocytes that already infiltrated the tumor in a given patient. As data in sarcoma indicates the immune system to be “broken a different way” in different patients, the PDE model could allow us tailor our therapeutic approach. Our results suggest that a heterogeneous T-cell population would indeed be beneficial in ACT for sarcoma (and other tumors).

Taken together, these approaches demonstrate a framework in which we first simplify a specific problem to make it tractable, in this case through dimensionality reduction using Boolean networks, and then tailor it to a specific question. Our ODE model is focusing on the dynamics and balance between lymphocyte sub-populations, enabling it to investigate scenarios in which drugs modulate the interactions between these actors, such as checkpoint inhibitors. The PDE model on the other hand focuses on tumor heterogeneity, allowing us to study situations in which exert more control over the T-cell phenotype, as is the case in adoptive cell transfer approaches.

Different disease sites might require differing approaches, as additional lymphocytes population play important roles or other factor come into play, though the framework described above could guide us in finding answers to a variety of open question in the modeling of immunotherapy.

### 5.1 Future work

To ensure clinical applicability, we will center our efforts on two clinical questions formulated as the following specific aims:

**Specific aim 1: Identify phenotypic signatures of ex vivo sarcoma-derived TIL that predict ACT efficacy.** Hypothesis: interactions between lymphocyte sub-types in the tumor microenvironment are preserved ex vivo and result in a 'meta-phenotype' that impacts clinical potency. Further, this meta-phenotype can be empirically characterized.

Objectives:

1. Perform more detailed phenotypic analyses to define the relative proportions of functionally distinct sub-populations of sarcoma-derived TIL. These analyses will include innate lymphocytes, which have direct killing capacity and immunoregulatory potential.
2. Using the recently established CD107 marker as a proxy for the potency of anti-tumor response within different TIL isolates, identify correlations between the relative presence of distinct lymphoid sub-populations and the resultant clinical response.

**Specific aim 2: Construct mathematical models to characterize the optimum patient-specific TIL meta-phenotype in metastatic sarcoma.** Hypothesis: Existing functional knowledge of TIL sub-populations in patients treated in clinical trials at Moffitt Cancer Center is sufficient for the development of mechanistic mathematical models that predict their dynamic interactions and clinical outcome following ACT.

Objectives:

1. Use a Boolean network approach to identify stable immunoregulatory motifs amongst lymphoid sub-populations. Incorporate these results in an ordinary differential equation model of tumor growth and immune interactions to predict the dynamic response to ACT.
2. Using a partial differential equation approach, extend tumor growth model to include explicit consideration of trait space, relatable to molecular data.
3. Probe coevolution of tumor and lymphoid populations.
4. Calibrate models against data generated in clinical trials and predict the patient-specific optimal lymphocyte heterogeneity and diversity to optimize ACT.

## 6 Conclusion

The integration of available knowledge into functional models of cancer immune reactivity can identify optimal network configurations and suggest novel ‘pressure points’ for therapeutic interventions. Our work aims to improve the clinical efficacy of ACT over existing, poorly predictive, proxy methods for evaluating antitumor response. Furthermore, identifying predictive signatures of TIL meta-phenotype will advance our understanding of the complex functional interactions that have the capacity to drive the immune response toward reactivity or suppression. While this project focuses on metastatic soft tissue sarcoma, the knowledge gained through this project will inform more effective approaches to other cancers using a similar modeling framework.

## Acknowledgments

Acknowledgment: We would like to thank the Moffitt Cancer Center as a whole, and the department of Integrated Mathematical Oncology in particular, for providing generous travel grants for participants as well as hosting this productive workshop.

